# Multisite Thalamic Recordings to Characterize Seizure Propagation in the Human Brain

**DOI:** 10.1101/2022.12.01.518775

**Authors:** Teresa Q. Wu, Neda Kaboodvand, Mike Veit, Ryan J. McGinn, Zachary Davey, Anjali Datta, Kevin D. Graber, Kimford J. Meador, Robert Fisher, Vivek Buch, Josef Parvizi

## Abstract

Neuromodulation of the anterior nuclei of the thalamus (ANT) has shown to be efficacious in patients with refractory focal epilepsy, but it is not uniformly effective. One important uncertainty is to what extent thalamic subregions other than the ANT are recruited earlier and more prominently in the propagation of seizures in patients with presumed temporal lobe epilepsy (TLE). To address this unknown, we studied 11 patients with clinical manifestations of TLE planned to undergo invasive stereo-encephalography (sEEG) monitoring. We extended cortical electrodes to reach thalamic nuclear subdivisions in the anterior (ANT), middle (mediodorsal) and or posterior (pulvinar) sites. This multisite thalamic sampling was without any adverse events. Intracranial EEG (iEEG) recordings confirmed seizure-onset in medial temporal lobe, insula, orbitofrontal and temporal neocortical sites – highlighting the importance of iEEG for more accurate localization of seizure foci. Visual review of EEGs documented early and prominent involvement of specific thalamic sites. Seizures originating from the same brain origin produced a stereotyped thalamic EEG signature. Visual review of EEGs, validated with single-pulse corticothalamic evoked potentials, documented early and prominent involvement of thalamic sites that would have not been predicted given the anatomy of seizure onset zones. Pulvinar was involved earlier and more prominently than other sampled nuclear subgroups in 60% of patients, even though all patients had a presumed diagnosis of TLE prior to invasive monitoring. Our findings document the feasibility and safety of multisite sampling from the human thalamus and suggest that the anatomy of thalamic involvement may not be entirely predictable on the basis of clinical information or lobar localization of seizures. Future clinical trials can establish whether offering more personalized targets for thalamic neuromodulation will lead to greater meaningful improvements in outcome.

## INTRODUCTION

Understanding the precise nature of corticothalamic interactions has become increasingly relevant since thalamic neuromodulation has shown recent promise in the treatment of patients with drug resistant epilepsies(Gadot *et al*., 2022). For instance, bilateral stimulation of the human thalamus in patients with drug resistant epilepsies have shown to reduce seizures with favorable long-term efficacy and safety profiles, but the response to this mode of treatment has shown marked variability with localization of seizure foci(Fisher *et al*., 2010; Salanova *et al*., 2021a). A recent study found significant improvement of seizures originating from posterior cortical regions in 3 patients when the posterior thalamic region (pulvinar) was the target of responsive neurostimulation(Burdette *et al*., 2021).

To date, our understanding of thalamic involvement in the early phase of seizure propagation in the human brain has been limited to only a few studies where single thalamic site within each individual has been explored. In a seminal work with TLE patients, Guye and colleagues obtained single-site thalamic recordings from the medial pulvinar group or the posterior part of the dorsomedian nucleus and documented thalamic involvement in the early phase of seizure propagation in ~86% of patients(Pizzo *et al*., 2021). This was in keeping with studies showing that the field potentials recorded from the rodent thalamus follow the cortical onset of seizures by <2 seconds(Yang *et al*., 2016). Romeo and colleagues studied three patients with a single additional depth electrode targeting the midline thalamus and confirmed that the thalamic recording site was recruited at varying points of seizure initiation ranging from 0 to 13 seconds(Romeo *et al*., 2019). Chaitanya and colleagues(Chaitanya *et al*., 2020) extended one single cortical electrode to reach either the ANT or the mediodorsal (MD) and centromedian (CM) nuclei and reported correct placement in the ~77% of ANT and 91% of MD/CM cases and their early recruitment during seizures. A different view was presented in a study (Osorio *et al*., 2015) showing thalamic ictal onset preceded scalp onset by about 0.5-2s with electrodes recording from either ANT (in 2 patients) or CM sites (one patient) using chronically implanted Deep Brain Stimulation (DBS) device. While these studies have unanimously documented the involvement of the thalamus during early phases of seizure propagation, they have highlighted the need for a systematic observation of multiple thalamic sites during seizure propagation in the human brain – at the individual patient level.

To address this gap of knowledge, we designed the current study to characterize the involvement of three subdivisions of the human thalamus (i.e., the anterior, mid, and posterior subdivisions) during seizure onset in patients with drug resistant focal TLE. Inspired by the methodology used in prior studies(Arthuis *et al*., 2009; Evangelista *et al*., 2015; Yang *et al*., 2016; Romeo *et al*., 2019; Chaitanya *et al*., 2020; Pizzo *et al*., 2021), we developed a novel surgical approach for multi-thalamic sampling that utilized an orthogonal trajectory extending frontal and temporal opercular or insular electrodes to the three specific thalamic nuclear subgroups desired on a per patient basis. Uniquely, we pioneered a single trajectory that traveled through the massa intermedia to combine recordings from bilateral medial thalamic nuclei into one electrode. Doing so enabled us to achieve bilateral coverage across three distinct sites of the thalamus with fewer penetrations and thus improving the safety of our approach. We employed conventional visual review of EEGs during ictal events and validated our findings using repeated single pulse electrical stimulation approach. Our observations confirm the reports of earlier studies and provide novel information about the mode of involvement of different thalamic subregions during seizure propagation in a group of patients with limbic semiology.

## METHODS

### Patient Selection

We recruited 11 patients who were considered clinically to have temporal lobe epilepsy (TLE) of uncertain laterality and precise anatomical origin. Per routine clinical protocols in our institution, and research procedures and consents approved by the Stanford IRB, these patients underwent bilateral stereo-EEG (sEEG) recordings. Prior to the time of sEEG recordings, all patients completed a comprehensive set of evaluations, including detailed clinical history, neurological examination, neuropsychological assessment, structural MRI, and scalp EEG monitoring. The majority of patients completed additional imaging and neurophysiological studies as needed for pre-surgical planning, including functional MRI for language mapping, FDG PET study, and high-density electrical source imaging.

### Anatomic electrode targets

Approximate locations and number of electrodes, along with their trajectories, were planned in a multidisciplinary surgical epilepsy conference with detailed review of presurgical data leading to the clinical hypotheses of most likely seizure onset zones (SOZ).

Thalamic monitoring was achieved through extension of electrodes covering planned cortical zones; hence no additional implantations were necessary. Thalamic recordings were derived from the most internal leads of the single multi-contact electrode clinically required to explore the superior temporal gyrus, posterior temporo-operculo-insular, or frontal regions. Thalamic subdivisions that were monitored included the anterior nuclei, mediodorsal, and pulvinar nuclei on either side. Given clinical limitations, the final number of thalamic electrodes ranged from 1-5 to monitor 1-6 thalamic nuclei. We did not implant in the centromedian thalamic nucleus in this series because its location was not within the trajectory of most other targets, and because it may be more involved with non-temporal lobe epilepsies(Velasco *et al*., 1987; Fisher *et al*., 1992; Velasco *et al*., 2006; Zillgitt *et al*., 2022)

### Electrode trajectory planning

High resolution T1, fast gray matter acquisition T1 inversion recovery (FGATIR), and T1 post-contrast imaging were used for planning. All patients required frontal and temporal opercular and/or insular coverage. Trajectories were planned to traverse in an orthogonal plane to capture cortical frontal or temporal operculum, insula, and then extended into specific nuclei of interest in the thalamus. We did not systematically target particular regions within the thalamic subdivision, but aimed for placement in ANT near the termination of the mammillothalamic tract and in the medial portion of pulvinar. The priority for all trajectories was safety and avoidance of middle cerebral sulcal and pial perforating blood vessels. At the highest density, we implemented a novel 5 electrode multi-thalamic sampling approach. The first trajectory extended from frontal or anterior temporal operculum to anterior insula to anterior thalamic nucleus. A second trajectory extended from posterior temporal operculum, temporoparietal junction, or supramarginal gyrus to the posterior insula to the pulvinar nucleus. These two trajectories were replicated bilaterally. The fifth and final electrode started in mid superior temporal gyrus or pars opercularis, traversed mid-insula, and extended to the mediodorsal nucleus and through the massa intermedia to terminate approximately 1 cm into the contralateral mediodorsal nucleus. In this highest density montage, we captured six thalamic regions (anterior, middle, and posterior nuclear groups bilaterally) with five electrodes that also had coverage of desired cortical and insular regions superficially. We used only reduced diameter (0.86mm) electrodes (Ad-Tech Medical, Oak Creek, WI) to help ensure minimal disruption to tissue. Trajectories were optimized by avoiding middle cerebral vessels as well as minimizing the distance traveled through the sylvian fissure. This was to minimize risk of deflection as the reduced diameter obturating stylet and electrodes passed through the two pial boundaries.

### Intraoperative workflow

The patients were brought to the operating room where general endotracheal anesthetic was induced. Five bone fiducials were placed. A volumetric intraoperative O-arm^®^ (Medtronic Inc, Minneapolis, MN) CT scan was obtained with the fiducials. The image data set was then merged with the preoperative CT and T1 pre and post contrast MRI scans. The patient was placed in a Leksell head holder and positioned supine. The ROSA™ robot (Zimmer Biomet, Warsaw, IN) was then attached to the Leksell adapter and registered to the patient’s head using the bone fiducials. Registration was accepted once <0.5mm accuracy was achieved. The head was then prepped in the usual fashion. For each percutaneous trajectory, the ROSA robot was positioned coaxially. A small vertical stab incision was made with a #15 blade. A 2.4mm drill bit was then introduced through the ROSA drill guide, and the drill guide lowered coaxially all the way down to the scalp. Once through the inner table of the skull, a bolt (bone anchor) was placed. A reduced diameter (0.8mm) obturating stylet was passed slowly to create the trajectory. This was a critical step due to the need for precise targeting and passing through sylvian fissure. Once the stylet was passed to depth, a reduced diameter (0.86mm) electrode was passed to target depth, inner stylet removed, and tightened into the bolt cap. The trans-massa thalamic trajectory was always performed first, to ensure highest degree of accuracy prior to the chance for subtle brain shift. At the end of the procedure, a 3-0 chromic gut buddy stitch was secured around each anchor bolt, to close the small stab incision upon bedside removal of the electrodes and bolts.

### Co-localization of electrodes

A thin cut CT-Head was obtained after electrode implantation to confirm absence of intracranial hemorrhage. Additionally, the CT images were co-registered to MRI data for verification of the trajectory. Precise electrode positioning in the FreeSurfer surface space, voxel space and MNI space were automatically extracted by iElVis toolbox(Groppe *et al*., 2017). T1-weighted MRI scan was used to generate 3D cortical volume and subcortical segmentation using recon-all command of Freesurfer v6.0.0(Fischl, 2012). The post-implant CT scan was aligned to the pre-implant MRI using the *flirt* from the Oxford Centre for Functional MRI of the Brain Software Library(Jenkinson and Smith, 2001; Jenkinson *et al*., 2012) or using *bbregister* from Freesurfer(Greve and Fischl, 2009) to get the best results. We then manually labeled each electrode on the T1-registered CT image using BioImage Suite(Papademetris *et al*., 2006). The electrode coordinates in the native anatomical space were carefully inspected for every single electrode contact and manually labeled by a neurologist and anatomist (J.P.) based on the individual brain’s morphology and landmarks.

### Thalamic parcellatiorr

Individual contact center of mass was defined in native T1 space for each subject. Center of mass of each contact was converted into a scalar X,Y,Z coordinate in MNI space. A widely used MNI thalamic atlas developed by our colleagues, Thalamus Optimized Multi Altas Segmentation (THOMAS)(Su *et al*., 2019), was utilized to parcellate the nuclear location of each contact within the thalamus. In order to do this, a 1mm cubic voxel region was created around each contact center of mass (contact neighborhood). For each contact neighborhood, the fraction of voxels that overlapped with each thalamic nucleus in the THOMAS atlas was calculated. Contact neighborhoods that fell solely within a single THOMAS nuclear mask would have a value of 1 for that specific nucleus, while contact neighborhoods that had no overlapping voxels with a given nuclear mask would have a value assigned to 0 for that specific nucleus. Thus, for every contact neighborhood anatomically inside the thalamus, the fractional overlap with each THOMAS nuclear mask was calculated. Then for each THOMAS thalamic nucleus, this within-subject fractional overlap was summed across subjects across all thalamic contacts to generate overall contact neighborhood fractional overlap values. Lastly, each THOMAS nucleus was segregated based on the most common trajectory used to obtain that nuclear coverage (between anterior versus mid versus posterior thalamic electrode trajectories).

### Intracranial Recording

Signals were collected from multiple contact depth electrodes with center-to-center contact spacing of 3 mm. Continuous electrocorticography signal was acquired with a digital Nihon-Kohden EEG machine at a sampling rate of 1000 Hz, in combination with continuous video recording. High frequency filter, time-constant, as well as voltage sensitivity settings were adjusted to optimize visual detection of high-frequency oscillations (typically at 300 Hz high-frequency filter, 0.001s time-constant, 10 μV sensitivity) and thalamic signals (best seen at 300 Hz, 0.1s, 10 μV). A sEEG bipolar montage including all channels was used for signal detection. Channels with excessive artifacts obscuring EEG signals were excluded from analysis.

### Identification of Ictal Patterns

All seizures captured were reviewed for onset zones. The cortical seizure onset zones were determined by visual analysis by the primary inpatient epilepsy team. The thalamic onset was determined by visual analysis of the sEEGs. Cortical and thalamic ictal onset signals were inclusive of various morphologies, such as pathologic high frequency oscillations, evolving fast activity, rhythmic spikes, or rhythmic spike-waves. Two epilepsy fellows and a senior attending, blinded to the diagnosed cortical seizure onset zones, reviewed the EEG tracings of seizures and answered three specific questions: A) Is the seizure propagated to the thalamus; B) which subregion of the thalamus is involved first; and C) which other thalamic sites are engaged next and in which temporal order. The raters were given randomly selected seizures from each individual patient. If a patient had more than one seizure type during their inpatient monitoring, we selected randomly 3 seizures from each seizure pattern. In selection of seizures from each individual patient, we relied on the clinical notes available in the electronic medical records prepared by the original clinical team caring for the patient during inpatient sEEG monitoring. Only 1-2 seizures were included if fewer than 3 seizures were captured for a specific seizure pattern. After completion of the ratings, we measured the interrater reliability of the three readers’ assessments of thalamic involvement and assessed 1) agreement on whether the thalamus was involved during ictal propagation, and 2) agreement on which thalamic nucleus was first involved at or following cortical onset. For the first assessment, Cohen’s Kappa was calculated based on the first two reviewers’ impressions. For the second assessment, consensus was determined with at least 2/3 agreement by independent review, as well as with group review and discussion.

### Study of cortico-thalamic evoked potentials (CTEPs)

To obtain an independent measure of effective connectivity between SOZs and each thalamic site we used the well-known method of repeated single electrical pulses as described before(Matsumoto *et al*., 2004). Single pulse stimulations (N=45) were performed with a bipolar setup using a cortical stimulator while the subjects were awake and resting. Single pulses of electrical current (5 mA, biphasic, 500 μs/phase) were injected between pairs of all adjacent intracranial electrodes at a frequency of 0.5 Hz (in total 90 seconds for each pair of electrodes). Electrical potentials were simultaneously measured in all other electrodes with a sampling rate of 1000 Hz. As we were interested in only the cortico-thalamic networks, electrodes in the white matter were excluded from the analysis. Specifically, we kept only electrodes which contained at least 1 grey matter voxel within a 2 mm sphere centered on the electrode. To minimize volume conduction effects, we also discarded data recorded from electrodes on the same electrode shaft as the stimulated electrode.

Two groups performed the analysis of evoked thalamic responses independently from each other to ensure validity of findings. The two approaches (i.e., CTEP Pipeline 1 and CTEP Pipeline 2) were generally similar but were different in some important details. The main differences of the two pipelines can be summarized as follows. CTEP Pipeline 1 included spectro-spatial decomposition spatial filtering to down-weight 1/f background nonperiodic activity, as well as removing electrical noise artifacts by fitting an autoregressive model minimizing the Akaike information criterion over the remaining samples, whereas the CTEP Pipeline 2 only considered notch filtering at powerline noise frequencies. Afterwards the signals were re-referenced to a bipolar montage in both pipelines, yet with different approach to segmentation. CTEP Pipeline 1 segmented the data into 2200 ms long trials that were time locked to the onset of the electrical pulses (200 ms pre-stimulus to 2000 ms post-stimulus), while the CTEP Pipeline 2 considered shorter 650 ms-long segments of the evoked responses starting from 150 ms pre-stimulus to 500 ms post-stimulus. In both methods, the time series data underwent L2-normalization, followed by normalization to the mean and standard deviation of the baseline activity (i.e., −200 to −20 ms in CTEP Pipeline 1 and −150 to −10 ms in CTEP Pipeline 2). Finally the signals were averaged over all trials of a given stimulation. Because the direction of activity is ambiguous in data collected from bipolar electrodes, we chose the sign of the time series so that the maximum evoked response was positive-as detailed in our(Veit *et al*., 2021) and others’(Keller *et al*.,2014) prior publications. Finally, peaks were detected for all recorded signals and their inversed versions, using MATLAB peak detection algorithm and a pipeline-specific thresholding procedure.

In the CTEP Pipeline 1, the distributions of the raw voltage and prominence of all peaks of the time-locked average voltage traces within the first 500 ms across all stimulation-recording pairs were defined for each individual. Afterwards, a two-level thresholding strategy was employed. First, a conservative threshold of average voltage/prominence trace plus 2 standard deviations was imposed to determine if a recording channel was activated in response to the electrical stimulation. On the second level, a more liberal thresholding of average voltage/prominence trace exceeding 1 standard deviation was used to detect the significant peaks for the channels which were identified at the first level. In the CTEP Pipeline 2, the minimum prominence was set as 7 times the standard deviation of the mean pre-stimulus baseline for each time series. Thalamic nuclei channels which did not show a significant response were discarded from the analysis. The order of occurrence and prominence of the post-stimulus peaks were compared between different thalamic nuclei. We were specifically interested in the latency of the first evoked peak in the recordings from the thalamus.

Pipeline 1 identified the earliest responding thalamic channel in each seizure pattern; Pipeline 2 identified the thalamic channel that has the earliest and most prominent response in each seizure pattern.

## RESULTS

### Patient Characteristics

11 patients (7 males and 4 females) were included in this study, with age range from 19-52 years. Mean IQ percentile was 66.7 (range 23-88, SD 19.2). Mean epilepsy duration was 14 years (range 2-35, SD 10.2). Six cases were non-lesional. For the 5 lesional cases, structural and pathological findings revealed a ganglioglioma (WHO Grade 1), gray matter heterotopia, focal dysplasia vs. prior injury, focal herniation through a skull base defect, and nonspecific reactive changes (**Table 1**). Mean number of electrodes placed per patient was 15 (range 10-21, SD 3.5), which included contacts extending to monitor 1-6 thalamic subdivisions (anterior, mid, and posterior on each side). Detailed pre-surgical data are summarized in **Table 2.**

**Table 1.** Patient characteristics.

| Patient # | age | gender | handedness | IQ Percentile | Epilepsy Duration (yrs) | Lesional (Y/N) | Pathology |
| --- | --- | --- | --- | --- | --- | --- | --- |
| 1 | 37 | F | R | 82 | 2 | N | n/a |
| 2 | 23 | F | L | 55 | 3 | N | n/a |
| 3 | 23 | M | R | 23 | 19 | Y | Ganglioglioma, WHO Grade 1 |
| 4 | 40 | M | R | 75 | 24 | N | n/a |
| 5 | 52 | M | R | 82 | 35 | N | n/a |
| 6 | 47 | M | R | -- | 17 | Y | Nonspecific reactive changes |
| 7 | 31 | F | R | 79 | 8 | N | n/a |
| 8 | 29 | M | R | 75 | 19 | Y | Gray matter heterotopia |
| 9 | 28 | M | R | 88 | 19 | Y | Focal dysplasia vs prior injury |
| 10 | 34 | F | R | 47 | 3 | N* | n/a |
| 11 | 19 | M | R | 61 | 4 | N** | n/a |

\*There was a L temporal encephalocele, which was not seizure focus
\*\*There were gray matter periventricular nodular heterotopia present, which were not seizure foci.

**Table 2.**
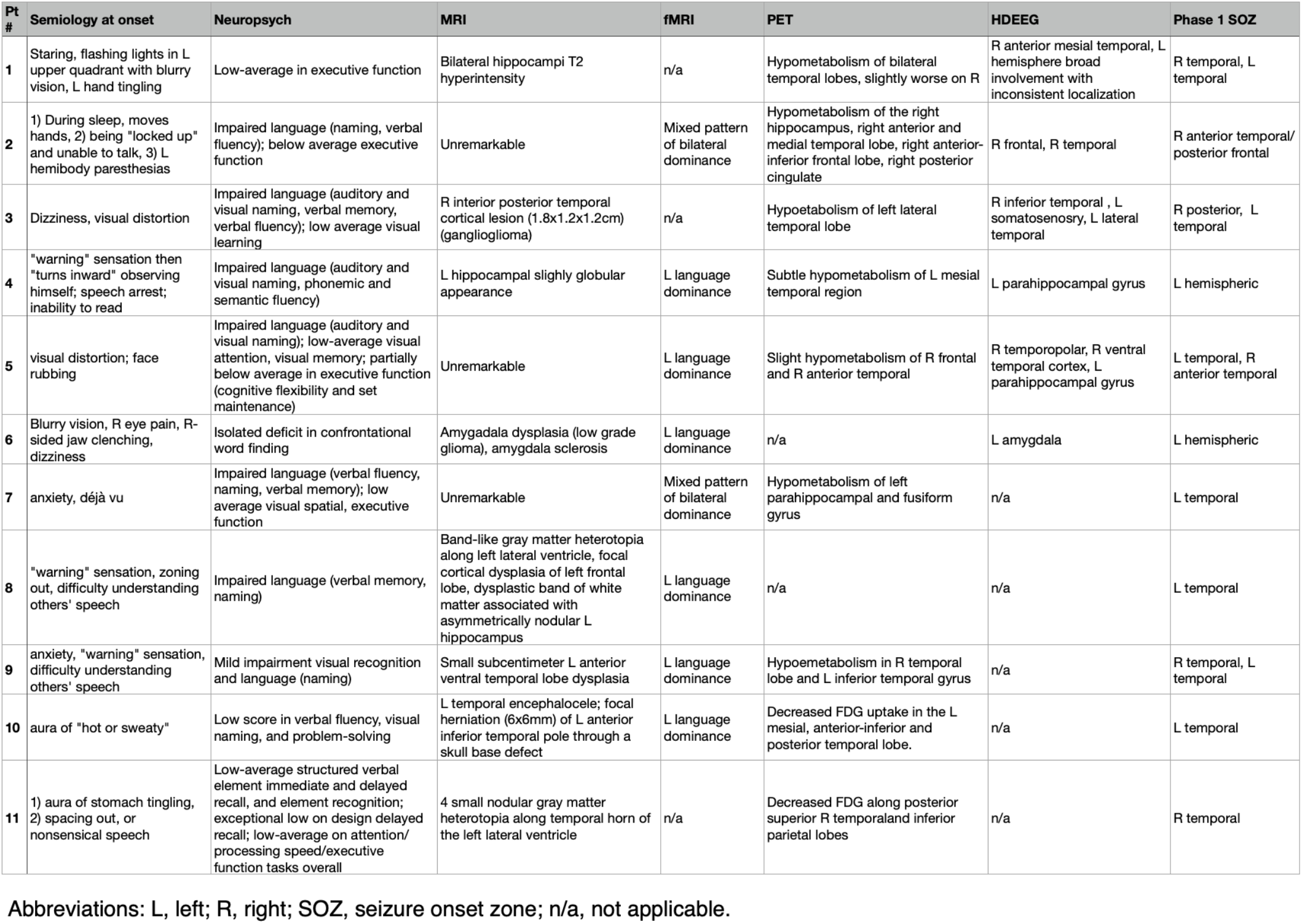
Seizure semiology and pre-surgical work up completed prior to Phase 2 intracranial monitoring.

### Anatomical Coverage

**Figure 2** demonstrates the MNI space locations of the electrodes, including overall 53 channels in the SOZ areas, as well as 33, 24 and 30 channels in the anterior, mid, and posterior thalamic regions. It is worth mentioning that all electrode locations have been superimposed in the right hemisphere, only for the visualization.

**Figure 1:**
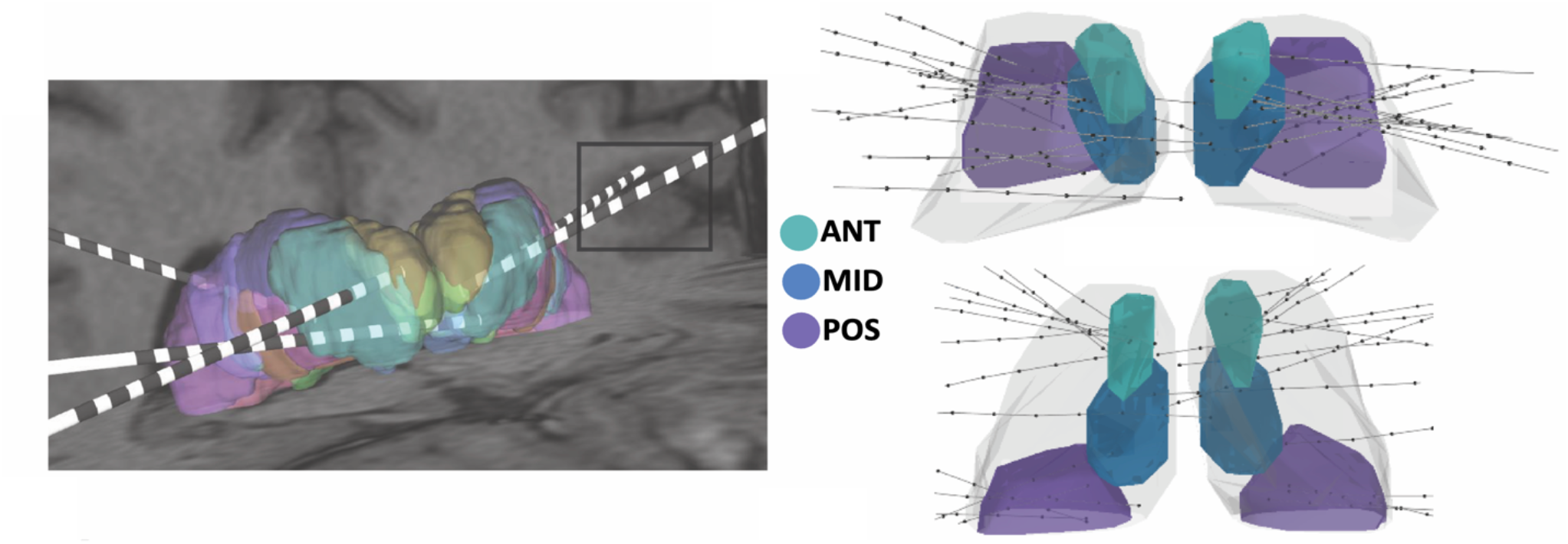
Thalamic Electrode Parcellation Across Subjects. A) Sample subject 3-dimensional reconstruction of a 5-lead orthogonal approach multi-thalamic sampling strategy. Includes bilateral anterior, posterior, and a single mid-thalamic trajectory that spans both thalami through the massa intermedia. The trajectories pass through opercular and insular structures which were desired by the clinical sampling strategy (black box) and are simply extended into thalamus. B) Axial and coronal views of an MNI space representation of all thalamic electrode trajectories across 10 subjects (lines) relative to the thalamus (gray). The AV nucleus of the ANT (cyan), MD nucleus (blue), and pulvinar nucleus (purple) from the THOMAS atlas are highlighted within the thalamic shape model. Individual contact center of mass is represented by black dots. C) Bar plot demonstrating the overall coverage of thalamic nuclei across all subjects and all trajectories. Nuclei were segregated based on the most common trajectory used to obtain that nuclear coverage (anterior trajectory = blue; mid trajectory = green; posterior trajectory = maroon). Target nuclei are highlighted in bold on the x-axis. D) Thalamic coverage only from either contact 1 (solid color) or contact 2 (patterned color) of each trajectory to analyze the contact neighborhood overlap of medial target nuclei in isolation.

**Figure 2:**
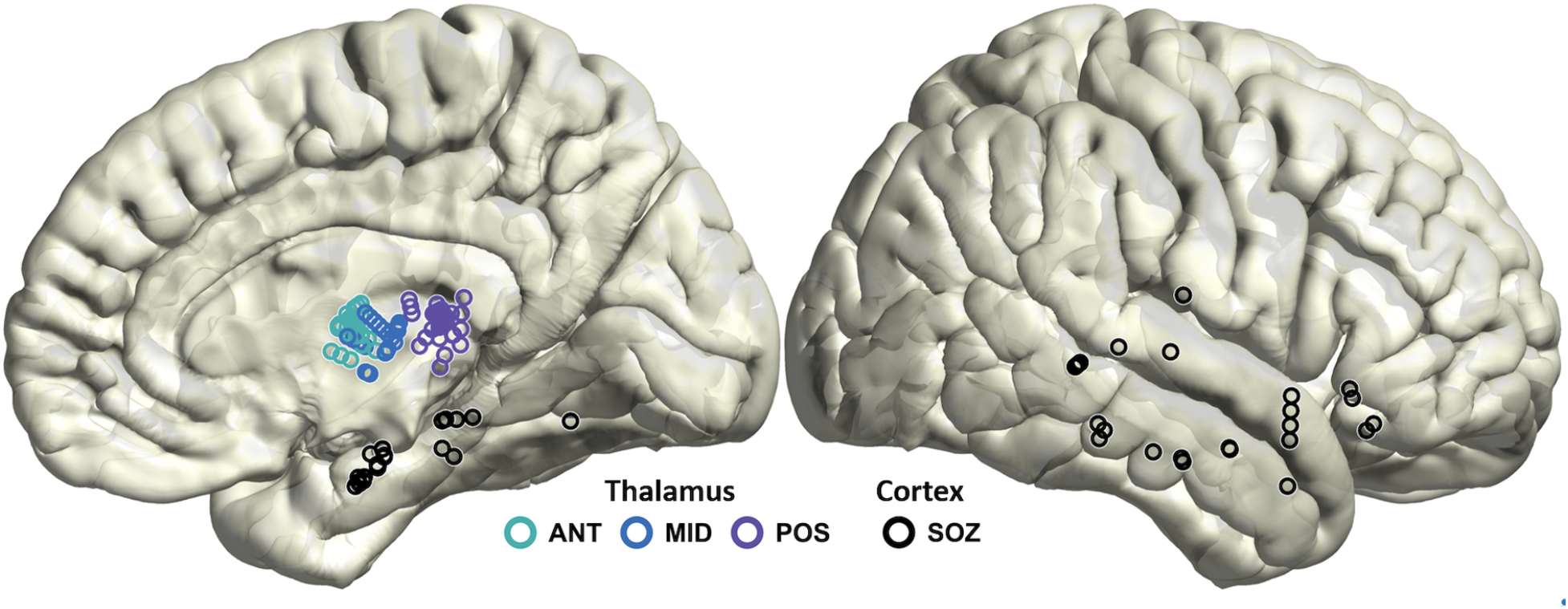
Electrode Coverage: Electrodes in the thalamus and seizure onset zones (SOZs) for all recorded seizures are shown on the right hemisphere. See Table 3 for additional details about SOZs. Each patient was implanted with many more electrodes throughout the brain, not shown here. Also, note that the electrodes are projected to the surface for 3D visualization purposes.

The pattern of thalamic coverage was not identical across all subjects because the thalamic electrodes were simply meant to be extensions of the appropriate cortical electrodes. **Table 3** shows the details of thalamic coverage in each case. In 9 out of 11 patients more than one thalamic nuclear subdivision was interrogated. In 7 of these 9 patients both ANT and pulvinar territories were covered simultaneously. In 8 out of 11 patients we were able to perform the CTEP procedure.

**Table 3.** Results from Phase 2 intracranial monitoring for seizure localization.

| Pt # | Phase 2 date | # of electrodes | Thalamic nuclei monitored | Sz Pattern | # of Sz | Seizure onset zone | First thalamic nucleus |
| --- | --- | --- | --- | --- | --- | --- | --- |
| 1 | 7/2/21 | 18 | R: ANT | A | 2 | R anterior hipp | R ANT |
|  |  |  |  | B | 8 | R middle temporal gyrus | R ANT |
|  |  |  |  | C | 2 | L orbital frontal | n/a |
| 2 | 7/23/21 | 13 | R: ANT, Pul | A | 6 | R posterior insula | R PUL |
| 3 | 8/28/21 | 17 | R: ANT, MD, Pul<br>L: ANT, MD | A | 21 | R peri-lesional, fusiform/inferior occipital gyrus | R ANT |
| 4 | 10/4/21 | 10 | R: MD<br>L: ANT, MD, Pul | A | 4 | L collateral sulcus, inferior temporal sulcus | L PUL |
|  |  |  |  | B | 4 | L inferior temporal sulcus, broad lateral temporal | L PUL |
| 5 | 10/19/21 | 21 | R: ANT, MD, Pul<br>L: ANT, MD, Pul | A | 13 | R orbital frontal + widespread R frontal and temporal | R PUL |
|  |  |  |  | B | 15 | L collateral sulcus | L ANT+PUL simultaneous |
| 6 | 12/6/21 | 14 | R: MD<br>L: ANT, MD | A | 17 | L temporal pole, anterior insula, anterior hipp, posterior hipp | L MD |
| 7 | 12/30/21 | 12 | R: MD<br>L: ANT, MD, Pul | A | 1 | L anterior hipp | L MD |
|  |  |  |  | B | 1 | L amygdala | L PUL |
| 8 | 1/28/22 | 11 | L: MD, Pul | A | 15 | L periventricular heterotopia - posterior portion of temporal horn of lateral ventricle | L PUL |
|  |  |  |  | B | 2 | L insula + transverse temporal gyrus, supra marginal gyrus | L PUL |
|  |  |  |  | C | 1 | L anterior hipp, L hipp heterotopia | L PUL |
|  |  |  |  | D | 1 | L periventricular heterotopia- frontal horn of lateral ventricle | n/a |
| 9 | 2/18/22 | 20 | R: ANT, MD, Pul<br>L: ANT, MD, Pul | A | 15 | L anterior collateral sulcus | L PUL |
|  |  |  |  | B | 1 | R PHG, mid hippocampal | R ANT |
| 10 | 3/4/22 | 13 | L: ANT | A | 11 | L anterior hipp | L ANT |
|  |  |  |  | B | 1 | L anterior, mid, posterior hipp | L ANT |
| 11 | 4/4/22 | 14 | R: ANT, Pul<br>L: ANT, Pul | A | 1 | L anterior hipp | L ANT |
|  |  |  |  | B | 3 | R anterior, posterior hipp | R ANT |

Included are the total number of electrodes implanted per patient, thalamic subregions monitored per patient, different seizure patterns captured from each patient with the number of seizures per pattern, the cortical seizure onset zone and first thalamic nucleus involved as determined by visual analysis. Abbreviations: L, left; R, right; Pt, patient; Sz, seizures; ANT, anterior thalamic nucleus; MD, mediodorsal thalamic nucleus; PUL, pulvinar thalamic nucleus; hipp, hippocampus; PHG, parahippocampal gyrus.

### Safety of Thalamic Implantation

There were no complications during or after implantation. There were no hemorrhages identified on the post-operative CT and no other complications were noted. All patients woke up neurologically intact (or at their preoperative baseline) following the implantation procedure. During the epilepsy monitoring unit (EMU) admission, all patients were able to participate fully in the clinical testing that was required. After explantation, minor pneumocephalus was a uniform finding across all patients. There were no clinical complications that occurred due to explantation. Patients were able to be discharged within 12-24 hours of explantation as desired.

### Seizure Onset Zones

A total of 145 seizures were captured and visually analyzed (2-28 seizures per patient). The seizures were grouped into 22 distinct patterns. Within each pattern, the seizures had the same onset zone and propagation pathway based on electrode sampling. Of the seizure patterns, cortical onset was noted to be on the left in 15, and on the right in 7. Six seizure patterns (from 5 patients), which consisted of 41 captured seizures, had broad regions of onset (> 1 identified cortical origin), where the exact onset location is unclear. The cortical onset zones were predominantly of temporal origin, including orbitofrontal, amygdalar, hippocampal, parahippocampal, insular, inferior temporal, and temporoparietal regions (**Table 3**).

### Thalamic Propagation of Seizures based on Visual EEG Review

There was a substantial agreement on whether the monitored thalamic nuclei were involved during seizures (percentage agreement was 98.0, Cohen’s Kappa between reviewers 1 and 2 was 0.79). Subsequently, to determine the level of agreement on which thalamic nucleus was first to be involved at seizure onset, a total of 37 seizures with more than one thalamic site monitored were included. In 34/37 seizures (92.0%), there was at least 2/3 majority agreement. In the remaining 3/37 seizures (all of which belonged to the same seizure pattern), one reviewer called the ANT to be first involved, one called the pulvinar, and the third called the ANT and pulvinar to be simultaneously affected. A group review was then conducted, with consensus achieved for anterior and posterior simultaneous involvement (see Methods for details). The thalamic onset with majority agreement was used for subsequent analysis. **Table 3** lists the first thalamic nucleus involved in each seizure pattern identified by the visual analysis.

### CTEP and Visual Review Comparisons

A summary of the results is provided in **Table 4**, which includes data from 8 patients (total of 13 seizure patterns), with relevant cortico-thalamic evoked potential data. Subsequent comparisons of the CTEP-identified thalamic response using two independent pipelines and the visual analysis method revealed agreement in 11/13 seizure patterns. There were two seizure patterns with discrepant results. The first case (patient #1, seizure pattern C) had the cortical seizure onset zone in the left hemisphere, while the implanted thalamic electrode was on the right – both CTEP Pipeline 1 and visual review did not appreciate an early thalamic signal, whereas Pipeline 2 detected signal in the right anterior thalamus, albeit at prolonged latency of >200ms. In the second case (patient #3, seizure pattern A), the cortical seizure onset zone was in the right peri-lesional inferior posterior temporal region; however, CTEP Pipeline 1 did not appreciate a significant thalamic signal, whereas Pipeline 2 detected response in the posterior thalamus and visual review in the anterior thalamus. It should be noted that in the patient (patient #6) with CTEP recording who was not included in the table, there were no sufficient electrode contact pairs in the thalamic nucleus of interest to produce accurate CTEP comparisons. In this patient, visual analysis identified left MD to be the first thalamic nucleus involved. However, there was only one channel contact in the left MD (the pair of contacts spread between left and right MD nuclei) Hence according to this CTEP analysis, the earliest peaks were detected in the left ANT and bilateral MD. Additionally, in patient #3 with SOZ in the inferior temporal cortex, based on Pipeline 1 approach, the injected current to the lesion cavity did not induce any evoked potential in the recorded channels in the thalamic nuclei. According to the data from eight patients with valid relevant cortico-thalamic evoked potential data, the earliest significant peak in the recorded voltage occurred in the thalamic divisions which were initially identified as the first thalamic nuclei involved during a seizure (**Table 4**).

**Table 4.** Summary of cortico-thalamic evoked potential mapping.

| Pt # | Sz pattern | Cortico-thalamic evoked potential mapping |  |  |  | Visual method |
| --- | --- | --- | --- | --- | --- | --- |
|  |  | Seed |  | Thalamic response (Channel name) |  | Thalamic response (Channel name) |
|  |  | Anatomical label | Channel name | Pipeline 1 | Pipeline 2 |  |
| 1 | A | R anterior hippocampus | RMHP 3-4 | R ANT (RAI2-3) | R ANT (RAI1-2) | R ANT (RAI1) |
|  | B | R middle temporal gyrus | RMHP 5-6 | R ANT (RAI2-3) | R ANT (RAI1-2) | R ANT (RAI1-3) |
|  | C | L orbitofrontal | LORF 9-10 | – | R ANT (RAI1-2) | – |
| 2 | A | R posterior insula | RF 7-8 | R PUL (RH1-2) | R PUL (RH1-2) | R PUL (RH1-2) |
| 3 | A | R peri-lesioned | RD 3-4 | – | R PUL (RK2-3) | R ANT (RI1-2) |
| 4 | A | L inferior temporal sulcus | PHGa 3-4 | L PUL (PTH1-2) | L PUL + L MD (PTH1-2, MTH3-4) | L PUL (PTH1-2) |
|  | B | L inferior temporal sulcus | PHGa 3-4 | L PUL (PTH1-2) | L PUL + L MD (PTH1-2, MTH3-4) | L PUL (PTH1-2) |
| 5 | A | Orbitofrontal cortex | ROH 9-10 | R PUL (RPT3-4) | R PUL (RPT3-4) | R PUL (RPT2) |
|  | B* | – | – | – | – | – |
| 9 | A | L anterior collateral sulcus | LMPG 5-6 | L PUL (LpTH1-2) | L PUL (LpTH1-2) | L PUL (LpTH1-2) |
|  | B* | – | – | – | – | – |
| 10 | A | Anterior hippocampus | aHIP 1-2 | L ANT (LaTH2-3) | L ANT (LaTH1-2, 3-4) | L ANT (LaTh1) |
|  | B | Posterior hippocampus | pHIP 1-2 | L ANT (LaTH1-2) | L ANT (LaTH1-2) | L ANT (LaTh1) |
| 11 | A | L anterior hippocampus | LaH 1-2 | L ANT (LANT2-3) | L ANT (LANT1-2) | L ANT (LANT1) |
|  | B | R posterior hippocampus | RpH 1-2 | R ANT (RANT2-3) | R ANT (RANT1-2) | R ANT (RANT1) |

Included are comparisons of results from two independent analyses, from Pipeline 1 and Pipeline 2, as well as results from visual method. Highlighted are two seizure patterns with discrepancy between the three results. Abbreviations: L, left; R, right; Pt, patient; Sz, seizures; ANT, anterior thalamic nucleus; MD, mediodorsal thalamic nucleus; PUL, pulvinar thalamic nucleus.
Note: \*no cortico-thalamic evoked potential data was available.

### Cortical and Thalamic Onset Comparisons

Of all the seizures captured, ictal propagation was visually detected in the thalamus, following cortical onset, in 20/22 seizure patterns. None of the seizure started in the thalamus prior to cortical onset. In the 20 seizure patterns with thalamic propagation, the earliest thalamic signals occurred ipsilateral to cortical ictal onset zones. In the 2 seizure patterns without visually detected thalamic spread, one had a cortical onset zone in the contralateral hemisphere from the implanted thalamic electrodes, and the seizures remained focal within the contralateral hemisphere; one began and remained distinctly in gray matter heterotopia without further spread.

We next explored the relationship between the anatomy of the cortical seizure onset location and the subregion of the thalamus to first become involved. This was examined only in seizures with detected thalamic spread and with more than one thalamic nuclei monitored. Seizures originating from a periventricular heterotopia also were excluded. 15 seizure patterns with a total of 105 captured seizures met this criteria. Of these, 6 seizure patterns with 41 seizures had broad regions of onset (>1 identified cortical origin), while 9 seizure patterns with 64 seizures had captured a narrow, potentially surgically targetable, cortical region of onset on intracranial monitoring (**Table 5**). In seizures with narrow or broad regions captured at onset, the cortical geography was not predictable of which thalamic subdivision, whether anterior, mid, or posterior, became first involved in seizure propagation. For instance, of 51 seizures with onset in the inferior temporal cortex, 21 propagated first to the anterior thalamus, 15 to the posterior thalamus, and 15 to the anterior and posterior thalamus simultaneously. Additionally, 23/41 (56.1%) of the seizures with broad regions of onset, which included frontal and temporal regions, propagated to the posterior thalamus first. Hence, despite their well-known networks, the anatomic location of cortical origins did not predict first thalamic subdivision involvement.

**Table 5.** Seizures with captured cortical onset zones (narrow vs. broad) and their earliest thalamic propagation.

| Narrow cortical onset zone | Group | # Sz (from # pts) | Thalamic onset |  |  |  |
| --- | --- | --- | --- | --- | --- | --- |
|  |  |  | ANT | MD | PUL | ANT + PUL |
| anterior hippocampus | hippocampus | 6 (3) | 4 | 1 | 1 |  |
| anterior + posterior hippocampus |  |  |  |  |  |  |
| amygdala | amygdala | 1 (1) |  |  | 1 |  |
| posterior insula | insula | 6 (1) |  |  | 6 |  |
| collateral sulcus | Inferior temporal cortex | 51 (3) | 21 |  | 15 | 15 |
| anterior collateral sulcus |  |  |  |  |  |  |
| Fusiform gyrus |  |  |  |  |  |  |
| Broad cortical onset zone |  | # Sz (from # pts) | ANT | MD | PUL | ANT + PUL |
| frontal and temporal |  | 13 (1) |  |  | 13 |  |
| hippocampal + PHG |  | 1 (1) | 1 |  |  |  |
| temporal + insular |  | 17 (1) |  | 17 |  |  |
| insular + supramarginal gyrus |  | 2 (1) |  |  |  | 2 |
| collateral sulcus + ITG |  | 4 (1) |  |  |  | 4 |
| ITS + broad LTL |  | 4 (1) |  |  |  | 4 |

Note only seizures with >1 thalamic regions monitored were included. Abbreviations: # Pt, number of patient; # Sz, number of seizures; ANT, anterior thalamic nucleus; MD, mediodorsal thalamic nucleus; PUL, pulvinar thalamic nucleus.

### Thalamic “IctalSignature”

While individual seizures within each seizure pattern carried a stereotyped electrographic morphology and cortical spread, thalamic ictal signal also appeared to be highly stereotyped. The first thalamic nucleus involved in every seizure within a seizure pattern remained consistent. Additionally, while the morphology of the thalamic ictal signal differed between seizure patterns, it remained nearly identical between seizures within a seizure pattern. **Figure 3** displays examples of stereotyped thalamic onset in two distinct seizure patterns.

**Figure 1:**
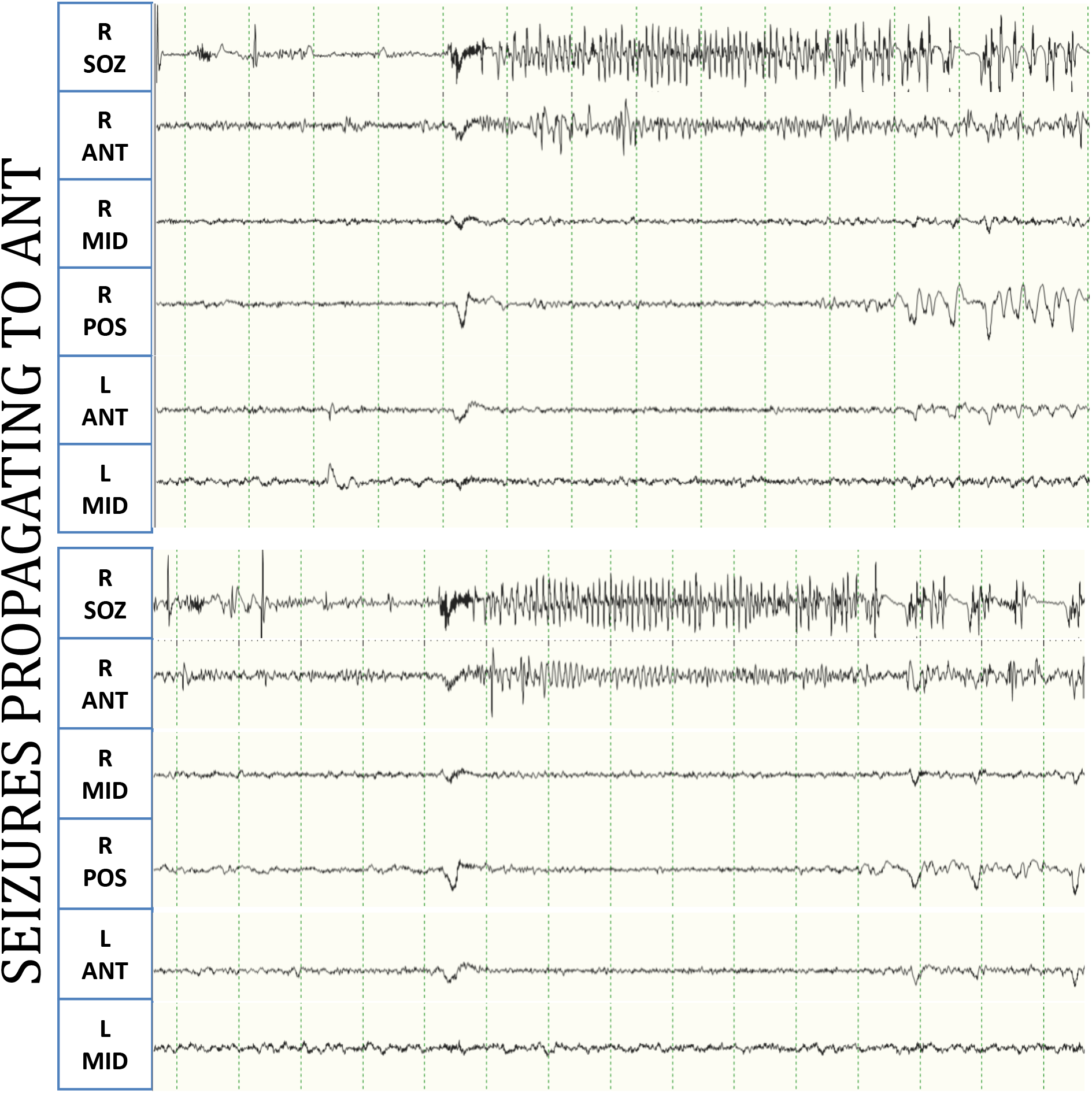
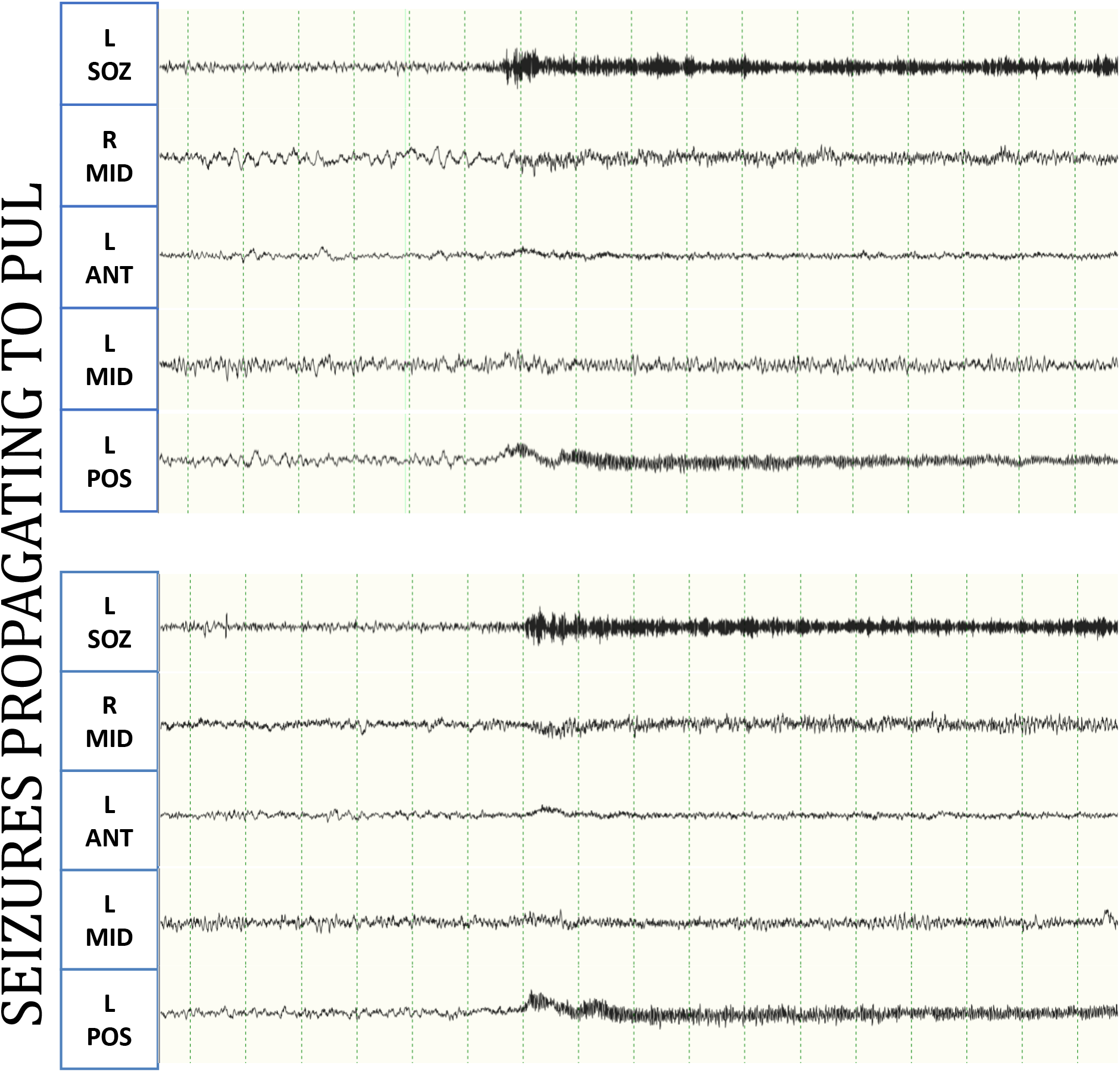
Thalamic ictal signal with stereotyped morphology, onset location and intra-thalamic propagation pattern. A) Two seizure examples from the same seizure pattern, with nearly identical early thalamic spread to the anterior thalamic nucleus. B) Two seizure examples from another seizure pattern, with nearly identical early thalamic spread to the pulvinar nucleus. Abbreviations: L, left; R, right; SOZ, cortical seizure onset zone; ANT, anterior thalamic subdivision; MID, mid thalamic subdivision; POS, posterior thalamic subdivision.

## DISCUSSION

We recruited 11 patients with presumed TLE and documented SOZs in not only the medial temporal lobe structures but also in the anterior and posterior insula, orbitofrontal lobe, and temporal neocortical sites, highlighting the importance of intracranial monitoring for increased accuracy of seizure origin. We used conventional visual review of EEG by clinicians and documented the involvement of different thalamic sites in most of our patients with multisite thalamic electrodes. In order to assess the validity of the visual review method in determining thalamic involvement, we measured the level of agreement amongst three independent clinician reviewers, as well as compared visual results with CTEP, using repeated single pulse electrical stimulation approach (Matsumoto *et al*., 2004). While a gold standard for determining thalamic subregion epileptogenicity is lacking, we observed substantial agreement in visual analysis amongst different reviewers, as well as between visual and CTEP analyses. Therefore, from a practical standpoint, we postulate that the expert visual review of the earliest and most salient epileptogenic site in the thalamus during a seizure can be used in the clinical setting to direct management strategies.

Our results show that different thalamic sites are involved differently in the very early stage of seizure propagation. This agrees with the extant literature, where thalamic participation in the epileptogenic networks of focal epilepsies has been documented (Ilyas *et al*., 2022; Piper *et al*.,2022). Pioneering past studies have shown increased synchrony between cortical and thalamic ictal activity in temporal lobe seizures, and that in a majority of cases the thalamic activity was driven by cortical activity (Guye *et al*., 2006). However, our findings with simultaneous recordings across multiple sites of the thalamus extend the existing evidence further and document important new findings that deserve attention and replication.

Our cohort included 11 patients who had been through several lines of presurgical diagnostic workup including 3T MRI, multiday inpatient observations with video EEG, and neuropsychological exams. On the basis of these diagnostic measures, the clinical diagnosis in majority of these patients was “likely temporal lobe epilepsy.” As such, all of these patients would have been candidates for ANT neuromodulation. However, as we report here, in a large proportion of these patients, nuclei other than ANT were engaged earlier and more prominently during seizures. Hence, our findings document that the earliest and most prominent thalamic involvement in a patient with presumed TLE could be of either the anterior, mid, or posterior thalamic subregions. This raises an important question of whether neuromodulation of the thalamus nuclei ought to be personalized to each patient.

Currently, stimulation of the ANT is proven to be a useful remedy for controlling seizures in patients with medication-resistant epilepsies (Fisher *et al*., 2010). However, the neuromodulation of ANT only benefits about two-thirds of patients (Salanova *et al*., 2015; Salanova *et al*., 2021b). It is presumed that patients with seizures originating from brain structures such as the hippocampus and medial temporal lobe - that are known from animal studies to communicate with and through ANT (Aggleton and O’Mara, 2022) to be the ones who benefit most from neuromodulation of this thalamic nucleus. Our findings, along with prior intracranial studies (Pizzo *et al*., 2021), suggests that the anatomy of thalamic involvement may not be entirely predictable based on the clinical semiology or lobar localization of seizures. In our study, location of SOZ was evidently not entirely predictive of which thalamic subdivision was going to be involved first and more prominently during seizures. For instance, some seizures of hippocampal, amygdalar, and insular origin were observed to first propagate to the pulvinar nucleus rather than ANT. Additionally, a notable number of seizures without a clear focal origin first propagated to the MD or pulvinar, including those with a broad frontal and temporal onset. This again highlights the possible additional clinical benefit of personalizing the target of thalamic neuromodulation based on intracranial monitoring.

It is worth highlighting that, similar to prior observations (Pizzo *et al*., 2021), we also noted a stereotyped “thalamic signature” in seizures of the same cortical origin. In other words, several seizures originating from the same cortical focus had near identical thalamic signature. In other words, in a patient with thalamic recordings, capturing different thalamic ictal footprint may indicate that that the patient’s seizures may be originating from different sources. In the future, and with additional evidence from larger number of patients, one may be able to compile an atlas of thalamic ictal footprints which will help identify the source of seizures or propagation pathways involving various brain regions. Additionally, an atlas of propagation pathways might usefully be correlated with data available from the Human Connectome Project on the other hand, if the treatment goal for a given patient is thalamic neuromodulation, one might argue that extensive cortical monitoring can be substituted by multisite thalamic recordings.

Using multisite thalamic recordings, one can develop personalized strategies for neuromodulation of individual thalamic nuclei in treating refractory focal seizures. Moreover, capturing information about the speed of first propagation to thalamic targets may provide valuable prognostic information as previously suggested *(Guye et al., 2006)*. Moreover, given that the thalamus contains distinct nuclei with distinct neuroanatomical connectivity profiles, it is reasonable to expect that seizures originating from distinct cortical regions would recruit different subregions of the thalamus. As such, knowing the exact thalamic circuitry involved in seizure propagation may yield clinically important information about the source of seizures and the networks of brain structures involved. For instance, a thalamic onset prior to observed onset in implanted cortical sites may indicate that the real SOZ is being missed and hence necessitate further implantation. Also studies in the future can elucidate if sampling from the thalamus with fewer electrodes may provide information about lateralization of seizures more accurately than many more cortical probes. Lastly, future studies are needed to determine if added information from thalamic multisite recordings can help optimize seizure localization and thus better surgical outcomes in cases where the SOZ is resected.

Our results provide a proof-of-concept evidence for feasibility and safety of multisite thalamic sampling in each individual. We implanted patients using an orthogonal approach for sampling frontal or temporal operculum, followed by anterior, mid, or posterior insula. With these trajectories, we extended them into various desired regions of the thalamus without adding to the total number of implanted electrodes. For bilateral mid-thalamic coverage, the single mid thalamic electrode trajectory was performed through a novel trans-massa intermedia approach, thus decreasing the overall surgical risk by obviating the need for one additional trans-sylvian electrode. Additionally, we used the RoSA robotic platform (Zimmer Biomet, Inc), a reduced diameter obturating stylet (AdTech) and reduced diameter electrodes (AdTech, Inc). The obturating stylet was a crucial step, since these trajectories went through various parts of the sylvian fissure where there were no vessels on preoperative imaging. With this surgical approach, we observed no complications, no thalamic hemorrhage or edema, and no neurological symptoms post operatively.

One of the meaningful findings of our study pertains to cases in which the epileptic network is found to be more distributed than initially hypothesized on the basis of pre-operative evidence. In these cases, a significant number of seizures propagated to the mid and posterior thalamic regions as the earliest and most prominent sites. This pattern of engagement might either indicate that the actual seizure onset zone was more posterior than hypothesized, or that the ictal onsets in these cases were simply broad. Either way, it is reasonable to hypothesize that MD or pulvinar nuclei rather than ANT neuromodulation in these patients may have produced better clinical benefits.

This study has several limitations. First, the seizures analyzed are from a small number of patients included during the study period, which may not be representative of the overall patient population undergoing surgical considerations. Second, the captured seizures in our study are of temporal onset, which do represent the most common seizure types but are not representative of all focal onset locations. Furthermore, sEEG studies with depth electrodes are inherently bound to sampling very limited regions in the brain, as ethical concerns limited electrode placements to only regions hypothesized to yield useful clinical information for the patient. Despite a clinician’s best planning for where to place electrodes based on comprehensive presurgical data, there remains the possibility of missing the actual cortical onset zone due to lack of sampling in that region. This may be the case in particular for the seizures with captured broad regions at onset, and in effect the complete seizure network may not be accurately characterized by the intracranial recording. Finally, it remains to be determined if an early propagation site necessarily qualifies as the optimal location for neuromodulation.

In closing, we argue that systematic thalamic monitoring may be helpful in elucidating the optimal thalamic target for “personalized neuromodulation” since knowing cortical seizure onset zone is not entirely predictive of which thalamic nucleus is most involved at onset. This may explain why neurostimulation is less effective in some cases. Additionally, with future studies, thalamic ictal patterns may help direct cortical SOZ localizations, narrowing the target regions for potential surgical interventions. Latency of seizure propagation to the thalamus and intra-thalamic spread patterns are also worth further investigations, as this may yield significant prognostic information for resective, ablative surgical, or neuromodulatory approaches in the treatment of refractory epilepsies.

